# Simplicity in numbers: Implications of diversity for the community Bioturbation Potential (BP_c_)

**DOI:** 10.1101/2020.04.21.053728

**Authors:** Andreas Neumann, Alexa Wrede

**Affiliations:** Helmholtz-Zentrum Geesthacht, Institut für Küstenforschung, Max-Planck-Straße 1, 21502 Geesthacht

## Abstract

The mobility of particles and solutes within aquatic sediment is substantially impacted by faunal bioturbation activities such as bioirrigation and sediment reworking. The non-quantitative community bioturbation potential (BP_c_) aims to estimate the potential of a given benthic community for bioturbation activities, based on the functional traits of the species within that community. We argue that in sufficiently diverse communities, the characteristics of individual species, expressed as traits scores, average each other out, and approach a universal constant. This enables to derive a simpler community bioturbation potential (BPs), which may be applied in cases when taxonomic expertise or trait data are not available. As a proof of concept, we evaluated a dataset of 84 benthos assemblages, and found a high degree of agreement of full BPc and simplified BPs.

## Introduction

Aquatic sediments provide reaction zones for very diverse reactions such as denitrification and methanogenesis. The first reduces eutrophication by mitigating anthropogenic N loads while the latter produces a potent greenhouse gas (Devol 2015, Evans et al. 2019). Since both reactions occur within the sediment, their long-term reaction rate is ultimately limited by transport mechanisms for the resupply of substrate. Generally, the balance between sequestration and recycling is controlled by the intensity of exchange between sediment and water column. Of all exchange modes, molecular diffusion is the least intense one, which is exceeded by porewater-advection and biotransport by several orders of magnitude (Hüttel et al. 2003, Ahmerkamp et al. 2017, Aller & Cochran 2019). While pressure-driven porewater advection and bedform migration dominate in permeable sediment (Ahmerkamp et al. 2017), fauna-sustained exchange processes such as bioturbation and bioirrigation dominate in impermeable sediment (Hüttel et al. 2003).

Flow-sustained benthic exchange is a physical process and can be described with a very limited number of parameters. However, fauna sustained exchange between sediment and water column relies on complex biological communities with high variability in taxonomic and functional composition. This complicates the interpretation of biogeochemical observations or the modelling of benthic processes.

The community bioturbation potential (BP_c_) is an attempt to grasp this complexity by estimating the potential of actual benthic communities to bioturbate with a single non-quantitative value (Solan et al. 2004). To do so BP_c_ includes abundance, biomass and trait scores, which parameterize mobility and feeding characteristics, of each species within the community. The BP_c_ is calculated by Equation 1

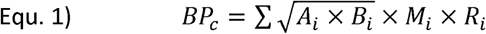

where *A*_*i*_ and *B*_*i*_ are abundance and biomass of species *i*, and *M*_*i*_ and *R*_*i*_ are the trait scores for mobility and reworking. Here, trait scores are integers in the range of 1 – 4 for *M*_*i*_, and in the range of 1 – 5 for *R*_*i*_, respectively. The trait product (*M*_*i*_ x *R*_*i*_) describes the contribution of abundance and biomass of a given species to the overall bioturbation. For the calculation of BP_c_ the benthos assemblage needs to be taxonomically determined, sorted, and weighted. Moreover, values of M and R need to be picked from look-up tables (e.g. Queirós et al. 2013) or determined by an expert by means of literature review or personal experience. The practical application of BP_c_ is thus dependent on the availability of experienced examiners and sufficient knowledge of each species to assign the values for M_i_ and R_i_. This in not an issue in well-studied regions such as the North Sea, but might become much more difficult for less covered regions.

In these cases, ecology might come to the rescue. Generally, species of a given habitat tend to reduce the competition for limited resources by reducing the overlap of their niches, which then results in a differentiation of traits (e.g. MacArthur & Levins 1967, Leimar et al. 2013). Competing species become dissimilar over time. As for the bioturbation potential, the specific trait products should thus be different among the species of a given benthos assemblage. This could imply that niche separation and differentiation of traits results in a balancing and compensation of individual traits when the community is sufficiently diverse. The trait-based community bioturbation potential BP_c_ of sufficiently diverse communities might thus become independent from the trait scores of the individual species and approach a universal, constant value.

If this hypothesis were true, then assigning a constant value for the trait product and measuring combined abundance and biomass of all species should provide a simplified bioturbation potential that is very similar to the corresponding values of the full BP_c_.

Unfortunately, the sum of species-wise calculated square roots is not equal to the square root of sums (2):

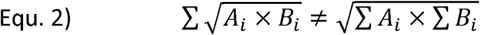

However, the error of equation 2 is a function of the species richness N (Fig. 1), and thus can be compensated with the term (0.5 N^-1^ + 0.5) (Equ. 3).

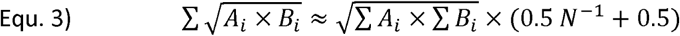

**Fig. 1:**
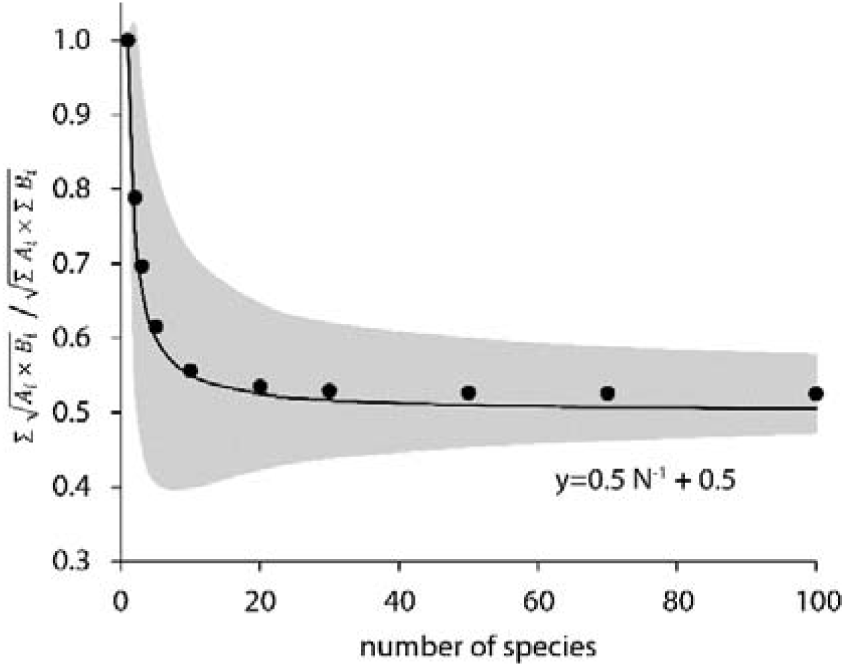
Average ratio of sum of roots vs. root of sums (full circles) as estimated with 10.000 random assemblages, 1 sd of averages is indicated by the grey band. The solid, black line indicates the approximation as applied in Equation 3.

The term (0.5 N^-0.9^ + 0.5) is approximately 0.5 for N > 10 (Fig. 1), and thereby Equ. 1 can be rewritten as

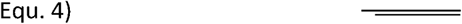

to estimate the effective trait score product (M x R)_eff_ for a given benthos assemblage. Lastly, a simplified community bioturbation potential (BPs) can be calculated as

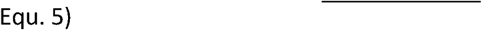

The scope of our present paper is to test the hypothesis that the full BPc can be replaced by a simplified BP_s_ for sufficiently diverse communities and to evaluate the applicability of the simplified BP_s_. We focus here entirely on the effect of species richness on the application of the BP_c_.

## Material & Methods

### Synthetic community data

Calculation of BP_c_ and (M x R)_eff_ for 10,000 random communities with 1 to 80 species based on randomly (Haahr 2019) assigned values for abundance, biomass, and trait scores.

### North Sea Intertidal benthos data

The data was derived from a monitoring program of the Lower Saxony Water Management, Coastal Defence and Nature Conservation Agency (NLWKN). The data covered 84 intertidal stations, sampled from 2006-2014 in 24 locations along the coast of Lower Saxony, Germany. Each station combined data of 10 replicated core (0.01 m^2^ cross sectional area) samples sieved through a 1 mm mesh and fixated in a 4% formaldehyde solution. Organisms have been determined to species level if possible and biomass was given as ash free dry mass. The dataset was taxonomically harmonized to prevent artificial diversity. The species spectrum was taxonomically reduced to cover only those species that accounted for 95% of both total biomass, abundance and frequency of occurrence (123 species). BPc trait scores were compiled from literature information and taken from Queirós et al. 2013.

## Results & Discussion

We established communities with 1 – 80 species and random values for abundance, biomass, and trait scores to estimate the effective trait product (M x R)_eff_ (Fig. 2 A). The average effective trait product (M x R)_eff_ is brought into perspective by comparison with the uncertainty of an individual trait product (M_i_ x R_i_). Although the traits M and R are established as categories, they are actually used as scalars to scale the effect of abundance and biomass on the resulting bioturbation. Since the values of these traits are discrete integers while the actual scaling is certainly rational, we argue that each trait score has an uncertainty in the range −0.5 to + 0.5 as the result of rounding to the next integer. Multiplying the two traits increases the uncertainty. In the case of M_i_ = 3 ± 0.5 and R_i_ = 4 ± 0.5, the true value of the individual trait product can be anywhere between 8.75 and 15.75. The uncertainty for other trait score combination is less than this rather extreme example and we estimated an average uncertainty of an individual trait product of 2.5.

**Fig. 2:**
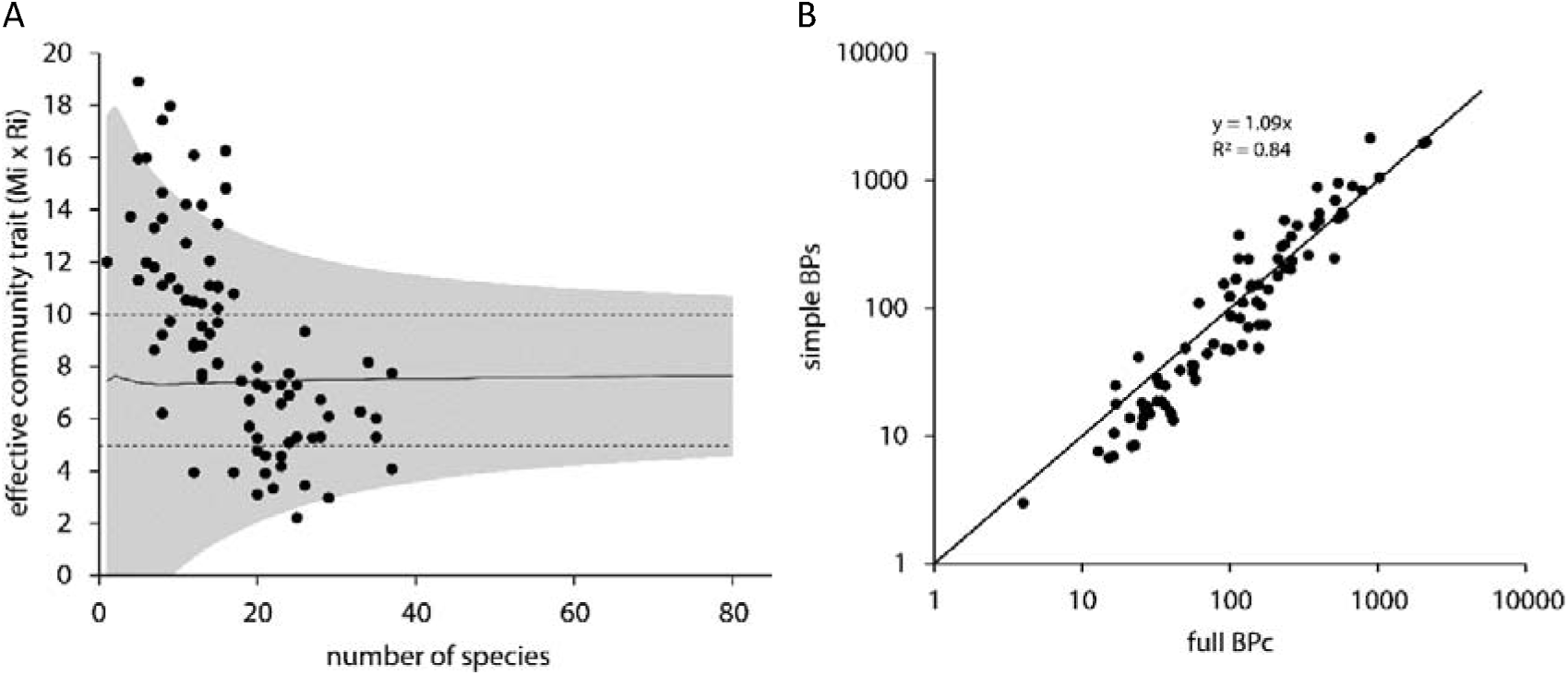
A: The averaged effective trait product (M x R)_eff_ and its uncertainty (2 standard deviations) as a function of the number of species in a random assemblage. The dashed lines indicate the uncertainty of an individual trait product (M_i_ x R_i_). Full circles indicate actual communities of the North Sea intertidal dataset (N = 84). See discussion for details. B: Simplified BPs (Equ. 5) vs. full BPc (Equ. 1) calculated for actual benthos assemblages of the North Sea intertidal dataset. The solid diagonal line indicates the 1:1 line (N = 84). See discussion for details.

Figure 2 A shows that the average effective trait product that was calculated with equation 3 for random communities is approximately 7.5, which is the expected value. The uncertainty of the effective trait product (Fig. 2 A, grey band) decreases asymptotically with the number of species and approaches the hypothetical uncertainty of a given trait product at a high species richness. For additional support, we evaluated an environmentally diverse dataset of 84 stations from the North Sea Intertidal. This data set comprised mud, fine sand, sand communities as well as a mussel bed and a coarse sand station. The effective trait products for 84 actual benthos assemblages from the North Sea Intertidal dataset calculated according Equation 4 agree well with the results from the random communities (Fig. 2). Starting from high values of the effective community trait product in the range of 12 to 18 in communities with low species richness, the values drop to 4 to 8 in communities with high species richness. Although the values of the effective community are within the 2 sd band that comprises 95 % of all cases, it appears that the effective community trait product of diverse communities is below the hypothetical value of 7.5 (Fig. 2 A). In our dataset, the lower values of the effective community trait product of samples with high species richness are the result of a significantly higher share of sessile species such as Mytilus, Balanus in the benthic community (p < 0.0001, Spearman r = 0.6816, N = 84). Some sessile species (e.g. Mytilus) may increase habitat complexity and provide hardsubtrate (Wehrmann et al. 2009) thus attracting further sessile commensals (e.g. Balanus) and enhancing biodiversity. These species do not contribute to the bioturbation of the sediment and have thus a very low trait product. However, the calculated values of the simplified BPs (Equ. 5) are well correlated (R^2^ = 0.84) with the values of the full community bioturbation potential BPc (Equ. 1).

From our present exercise, we can draw a few preliminary conclusions:

1. The BPs is not a replacement for BP_c_, especially not for communities with low species richness where uncertainties of results of equations 3 and 4 are high. However, it appears justified to apply the value 7.5 for the trait product (M_i_ x R_i_) in cases where values are unavailable and educated guess does not indicate otherwise (e.g. obviously sessile species).
2. Moreover, it appears even justified to apply the value 7.5 for all species of a sufficiently diverse community, which enables to estimate a simplified community bioturbation potential (Equ. 5). If this finding can be supported by results from other regions and additional datasets, our simplified BP_s_ may allow evaluation of benthic fluxes when abundance and biomass cannot be taxonomically resolved or traits may not be assigned due to a lack of literature data. In these cases, it would suffice to weight the unsorted fauna mass (biomass) and to count the number of heads (abundance) for the calculation of the simplified BP_s_. A question yet to be addressed is which communities are sufficiently diverse for the employment of the simplified BPs. Figure 2 A suggests that 50 or more species might be enough to let the 2 standard deviation interval (grey band) approach the uncertainty of an individual trait product (dashed lines). However, we found a high correlation of full BPc and simplified BPs in the North Sea Intertidal Dataset with a median species richness of just 15. And lastly, the proposed, simplified BPs aims for cases when the exact number of species of a given sample cannot be determined, and a threshold for the species richness is not very helpful in these cases. This implies that the simplified BPs demands at least the taxonomic experience to assess whether a given fauna assemblage is sufficiently diverse, and whether the general lifestyle mode does not indicate against BP_s_ application, for example a purely sessile community from a hard substrate.
3. Our conclusions should be verified with additional datasets. The analyses should also include the similar community bioirrigation potential (IP_c_), which was derived from the BPc (Wrede et al. 2018), and is different in details. A potential consequence could be that diverse benthos communities could have a similar ratio of bioirrigation potential and bioturbation potential although this is not necessarily true for the actual rates of bioirrigation and bioturbation. In this case, BPc and IPc would appear less suited to evaluate the individual contribution of each transport process.

## Acknowledgements

We are grateful for the North Sea Intertidal dataset that was kindly provided by the Lower Saxony Water Management, Coastal Defense, and Nature Conservation Agency (NLWKN).

## Notation

A: abundance (m^-2^)

B: biomass (Kg m^-2^)

BP_c_: full community bioturbation potential (Kg^0.5^ m^-2^)

BP_s_: simplified community bioturbation potential (Kg^0.5^ m^-2^)

M: mobility trait score (unitless)

N: number of species (unitless)

R: reworking trait score (unitless)

